# Baloxavir Marboxil Susceptibility of Influenza Viruses from the Asia-Pacific, 2012-2018

**DOI:** 10.1101/498766

**Authors:** Paulina Koszalka, Danielle Tilmanis, Merryn Roe, Dhanasekaran Vijaykrishna, Aeron C. Hurt

## Abstract

Baloxavir Marboxil (BXM) is an influenza polymerase inhibitor antiviral that binds to the endonuclease region in the PA subunit of influenza A and B viruses. To establish the baseline susceptibility of viruses circulating prior to licensure of BXM and to monitor for susceptibility post-BXM use, a cell culture-based focus reduction assay was developed to determine the susceptibility of 286 circulating seasonal influenza viruses, A(H1N1)pdm09, A(H3N2), B (Yamagata/Victoria) lineage viruses, including neuraminidase inhibitor (NAI) resistant viruses, to Baloxavir Acid (BXA), the active metabolic form of BXM. BXA was effective against all influenza subtypes tested with mean EC_50_ values (minimum-maximum) of 0.7 ± 0.5 nM (0.1–2.1 nM), 1.2 ± 0.6 nM (0.1-2.4), 7.2 ± 3.5 nM (0.7–14.8), and 5.8 ± 4.5 nM (1.8–15.5) obtained for A(H1N1)pdm09, A(H3N2), B(Victoria lineage), and B(Yamagata lineage) influenza viruses, respectively. Using reverse genetics, amino acid substitutions known to alter BXA susceptibility were introduced into the PA protein resulting in EC_50_ fold change increases that ranged from 2 to 65. Our study demonstrates that currently circulating viruses are susceptible to BXA and that the newly developed focus reduction assay is well suited to susceptibility monitoring in reference laboratories.

## 1. Introduction

Despite increased vaccination rates in many parts of the world, influenza continues to cause high levels of morbidity and mortality in high-risk groups [1], particularly when influenza seasons are dominated by A(H3N2) viruses [2]. Influenza antivirals are available for the short-term prophylaxis of individuals to prevent influenza infection, but their primary use has been to treat severely ill patients, many of whom are hospitalised. Two classes of influenza antivirals have been licensed for many years, the M2 ion channel inhibitors and neuraminidase inhibitors (NAIs). Clinical use of the M2 ion channel inhibitors, amantadine and rimantadine, is limited as close to 100% of circulating influenza A viruses contain an amino acid (AA) substitution at residue 31 of the M2 protein (S31N) that confers resistance to these compounds [3]. Four NAIs are licensed in different parts of the world, of which oseltamivir is the most widely available and commonly used. Oseltamivir resistance has been widespread amongst certain groups of viruses in different periods of time (e.g. seasonal H1N1 between 2007 and 2009 [4, 5]), but for the last seven years the frequency of viruses that circulate with reduced NAI susceptibility has remained at less than 5% [6]. The licensure of alternative antivirals, especially those with different modes of action to NAIs, is likely to be of benefit if oseltamivir resistant viruses emerge. In addition, combination therapy may be a strategy to improve clinical effectiveness compared to NAI monotherapy [7].

Baloxavir marboxil (S-033188, BXM) is an influenza polymerase inhibitor that was licensed for the treatment of uncomplicated influenza in Japan and the US in 2018 [8]. BXM is a prodrug that is hydrolysed by the enzyme arylacetamide deacetylase to the active form baloxavir acid (S-033447, BXA)[9]. BXA is a small molecule inhibitor of the highly conserved cap-dependant endonuclease (PA_N_) of influenza A and B viruses [10]. Inhibition of the PA_N_ disrupts endonuclease function and as a consequence the cap-snatching mechanism of the influenza polymerase [10]. Treatment of uncomplicated influenza with BXM as a single oral dose was shown in a Phase III clinical trial to reduce influenza symptom duration by 26.5 hours compared to placebo, similar to the reduction time achieved with oseltamivir [11]. However, 24 hours post-drug administration the reduction in viral load in BXM treated patients was twice as large as oseltamivir treated patients [11].

In post-treatment samples obtained from phase II and III clinical trials of BXM, A(H3N2) and A(H1N1)pdm09 viruses with reduced BXA susceptibility were detected and shown to carry AA substitutions at position 38 of the PA_N_ including I38T, I38M and I38F [12]. In the phase II clinical trial (which involved predominantly A(H1N1)pdm09 viruses) a substitution at I38 emerged in 4 of 182 patients (2.2%), while the phase III study (which involved predominantly A(H3N2) viruses) showed a frequency of an I38 variant in 36 of 370 (9.7%) of BXM recipients [11]. The highest frequency of viruses with reduced BXM susceptibility has been reported from a paediatric study, where 18 of 77 (23.4%) of patients had viruses bearing substitutions at I38 following treatment [12]. The transmissibility of the PA_N_ /I38 variants between patients in the absence of drug treatment is currently unknown, but *in vitro* studies suggest that these viruses do not replicate as well as the equivalent wildtype strains [10].

Given the frequency of viruses detected following BXM treatment with reduced BXA susceptibility, and the potential for widespread use of the drug in clinical practice, it is important to conduct surveillance of circulating strains for BXM susceptibility. This study aimed to develop a high-throughput and reproducible phenotypic assay for surveillance purposes to determine the BXA susceptibility of recently circulating influenza viruses in the Asia-Pacific region, thereby providing baseline susceptibility data to which prospective samples can be compared to in the future.

## 2. Materials and Methods

### 2.1 Antiviral Compounds, Cells and Viruses

20 mM stocks of Baloxavir acid (S-033447; BXA) (kindly provided by Shionogi & Co., Ltd.) were prepared in dimethyl sulfoxide (DMSO) (Sigma Aldrich, USA), filtered with a 0.2 μm surfactant-free cellulose acetate (SFCA) filter (ThermoFisher, USA) and stored in aliquots at -20°C.

COS-7 African green monkey kidney cells (ATCC, CRL-1651), HEK-293T (ATCC, CRL11268), Madin-Darby canine kidney (MDCK)-TMPRSS2 cells (kindly provided by Dr. Jesse Bloom) and MDCK-SIAT cells (MDCK cells that overexpress α2,6-linked sialic acids, kindly provided by Dr M. Matrosovich [13]), were cultured at 37°C in a 5% CO_2_ gassed incubator in Dulbecco’s Modified Eagle Medium (DMEM) (SAFC Biosciences, US). DMEM growth media (DMEM GM) used for the culture of COS-7, HEK-293T and MDCK-TMPRSS2 cell lines was supplemented with: 10% foetal bovine serum (Bovogen Biologicals, Australia), 1x GlutaMAX (Gibco, USA), 1x MEM non-essential amino acid solution (Gibco, USA), 0.06% sodium bicarbonate (Gibco, USA), 20 μM HEPES (Gibco, USA) and 100 U/mL penicillin-streptomycin solution (Gibco, USA). Growth media for MDCK-SIAT cells (DMEM GM SIAT) was supplemented as described for DMEM GM with the addition of 1 mg/mL Geneticin (Gibco, USA).

The influenza viruses used in this study were submitted through the WHO Global Influenza Surveillance and Response System (GISRS) to the WHO Collaborating Centre for Reference and Research on Influenza, Melbourne, Australia. Viruses tested were collected during 2013 to 2018 from Australia (n=158), Singapore (n=58), Malaysia (n=34), Cambodia (n=19), Thailand (n=7), Sri Lanka (n=3), New Zealand (n=3), New Caledonia (n=2) and Fiji (n=2). All viruses grown for assay purposes were propagated in MDCK-SIAT cells, using DMEM maintenance media (DMEM MM) supplemented as DMEM GM, with the addition of 4 μg/mL TPCK-treated trypsin (Worthington, USA) but without foetal bovine serum.

### 2.2 Site Directed Mutagenesis and the Generation of Recombinant Viruses

The reverse genetics plasmid pHW2000 (kindly provided by Richard Webby) containing each of the eight gene segments of A/Perth/261/2009 (A(H1N1)pdm09), A/Perth/16/2009 (A(H3N2)) or B/Yamanashi/166/98 virus (B/Yamagata lineage) were utilised for reverse genetics. The Gene Art^®^ Site-Directed Mutagenesis Kit (Life Technologies, USA) and relevant primer pairs were used for site-directed mutagenesis to introduce AA substitutions within the PA gene of each virus. Sanger sequencing was used to confirm AA changes in each plasmid. An eight-plasmid reverse genetics method adapted from Hoffman et al was used to generate recombinant viruses [13]. Alterations to the methods include, HEK-293T and MDCK-TMPRSS2 were seeded at an 8: 1 ratio with a total of 5x10^5^ cells and COS-7 and MDCK-TMPRSS2 at a 3:1 ratio with a total of 3x10^5^ cells, for influenza A and influenza B virus experiments, respectively. GeneJuice^®^ Transfection reagent (Merck Millipore, USA) was used for transfection. MDCK-TMPRSS2 were infected 72 hours post-transfection using 1 mL of supernatant from the co-culture. Virus growth was determined using haemagglutination assay and 1% (v/v) turkey red blood cells.

### 2.3 BXA Cytotoxicity Assay

The cytotoxicity of BXA was measured to identify non-toxic drug concentrations suitable for use *in vitro*. The inner wells of 96 well plates (Corning, USA) were seeded with MDCK-SIAT cells at a concentration of 2.5 x 10^5^ cells/mL (100 μl/well) and incubated overnight at 37°C in a 5% CO_2_ gassed incubator. The BXA concentration range of 50 μM to 0.4 μM was obtained from a serial two-fold dilution of BXA in DMSO (Sigma Aldrich, US) and further diluted in the final MM supplemented with 2 μg/mL TPCK-trypsin. One well was left free of BXA as a negative control. Treated cells were incubated at 35°C in a 5% CO_2_ gassed incubator for 24, 48 or 72 hours. Cell viability was determined using the CellTiter-Glo^®^ Luminescent Cell Viability Assay as per manufacturer’s instructions (Promega, USA) and luminescence was measured using a FLUOstar Optima luminometer (BMG Labtech, Germany). The BXA concentration that reduced cell viability by 50% compared to the cell only control (CC_50_) was calculated using non-linear regression analysis (GraphPad Prism, USA).

### 2.4 Virus Titration

Virus titration is required to select a suitable virus dilution for the focus reduction assay. MDCK-SIAT cells were seeded in the inner wells of 96 well plates (Corning, USA) at a concentration of 2.5 x 10^5^ cells/mL (100 μl/well) and incubated overnight at 37°C in a 5% CO_2_ gassed incubator. The experiment was only continued if the cell monolayer was 100% confluent the following day. MDCK-SIAT cells were infected and immunostained with previously described methods [14]. Briefly, nine half-log dilutions of viruses were prepared in MM. MDCK-SIAT cell monolayers were removed of SIAT-GM and washed once with PBS. 50 μl of each virus dilution was added to the appropriate wells on each plate with the tenth well mock-infected with MM to serve as a cell-only control. The plates were incubated at 35°C in a 5% CO_2_ gassed incubator for 90 minutes. The virus inoculum was then removed, cells washed once with PBS and overlayed with 100 μl of infection media (IM). IM contained equal parts 3.2% carboxymethyl cellulose (CMC) (1.6% final) (Sigma Aldrich, US) and 2x MEM (1x final) (Sigma Aldrich, USA) and was supplemented with 2 μg/mL trypsin. The 2x MEM was supplemented with 20 μM HEPES (Gibco, USA), 100 U/mL Penicillin-Streptomycin (Gibco, USA), 0.06% Sodium Bicarbonate (Gibco, USA). Plates were incubated at 35°C in a 5% CO_2_ gassed incubator for 24 hours. Following the incubation, the cells were fixed with 10% formalin (Sigma Aldrich, US) and permeabilised with 0.5% Triton X-100 (Sigma Aldrich, US). Plates were washed three times in wash buffer (0.05% Tween20 (Sigma-Aldrich, US) in PBS) and incubated for one hour with mouse anti-influenza monoclonal antibody against influenza A virus nucleoprotein (Millipore, USA, Cat#MAB8251) or influenza B virus nucleoprotein (Millipore, USA, Cat#MAB8661), diluted 1: 10,000 in 2% skim milk. Plates were then washed and incubated for one hour with goat anti-mouse IgG-horse radish peroxidase (Biorad, US) secondary antibody, diluted 1:1000 in 2% skim milk. Plates were again washed and then incubated for ten minutes in the dark with TrueBlue™ Peroxidase Substrate (KPL, US) followed by three washes with distilled water, the water was then removed, and plates were allowed to dry. Focus forming units (FFU) were quantified using the Immunospot BioSpot 5.1.36 (CenturyLink Inc, US).

### 2.5 BXA Focus Reduction Assay (FRA)

The concentration of BXA required for a 50% reduction in FFU (EC_50_) was used to determine susceptibility of influenza viruses to BXA. MDCK-SIATs were seeded and infected as described in section 2.4, however, virus was diluted such that there was 1000–2000 FFU/well, as previously determined by virus titration. Cell monolayers were washed with PBS and eight wells were overlaid with serial 4-fold dilutions of BXA (200–0.01 nM) in 100 μL IM. IM only was added to virus and cell control wells. Plates were incubated and immunostained as described in Section 2.4. Each virus was tested in duplicate wells, the foci were determined as an average of duplicate wells as described above. The EC_50_ was only calculated if the FFU count was between 500 and 2500 FFU in the virus control well. Using the mean FFU, the percent inhibition of FFU was calculated with the following formula:

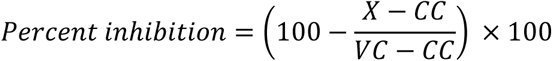

Where,

CC= FFU in cell control wells (no virus, no drug)

VC=FFU in virus control wells (virus, no drug).

X= Mean FFU

Using the percentage inhibition, the EC_50_ for BXA of each virus was determined using non-linear regression analysis (GraphPad Prism, USA).

### 2.6 Statistical Analysis

The Linear regression analysis and unpaired student’s t-tests were performed using GraphPad Prism (USA) where p-values < 0.05 were considered statistically significant. To evaluate assay reproducibility, FRA was performed with replicate (n = 48) wells of positive and negative controls (± virus) on a 96-well plate and Z factors were calculated using the equation outlined in [15].

## 3. Results

### 3.1 Cytotoxicity of BXA

To determine the maximum working drug concentration for use in *in vitro* assays, a CellTiterGlo assay was used to measure cell viability in the presence of increasing concentrations of BXA in MDCK-SIAT cells at 24, 48 and 72 hours. The MDCK-SIAT cell cytotoxicity (CC_50_-the 50% reduction of cell cytotoxicity compared to a cell only control) of BXA was 34.1 ± 1.9 μM, 10.1 ± 2.1 μM and 7.8 ± 0.9 μM at 24, 48 and 72 hours, respectively.

### 3.2 Reproducibility of BXA Focus Reduction Assay

Several factors were tested to ensure the reproducibility of the FRA method and it suitability for use as a robust, high throughput screening assay. Z scores were determined at 18, 24 and 30-hour post-infection times for three viruses, A/Perth/169/2017(A(H1N1pdm09)), A/Victoria/189/2017(A(H3N2)) and B/Sydney/42/2016 (B/Victoria lineage). The closer a Z score is to the value 1, the more reproducible the assay. For all influenza viruses the exclusion of the outer wells increased the Z-score by 10–30% and therefore was used for subsequent assays. For a 24-hour infection time, the Z score was 0.62, 0.73 and 0.69 for A(H1N1pdm09), A(H3N2) and influenza B test viruses, respectively. For influenza B, the Z score was similar at all infection times (0.58 to 0.69). In addition to these assays, two control viruses, A/Perth/16/2009 and A/Perth/16/2009-PA/I38M (reduced BXA susceptibility), were tested in (n=12) in biological replicates in distinct assays. The minimum and maximum EC_50_ values and the coefficient of variation was 0.13–1.19 nM (80%) and 6.06–12.49 nM (45%), respectively. The EC_50_ obtained in each experimental repeat is shown in Figure 1.

**Figure 1.**
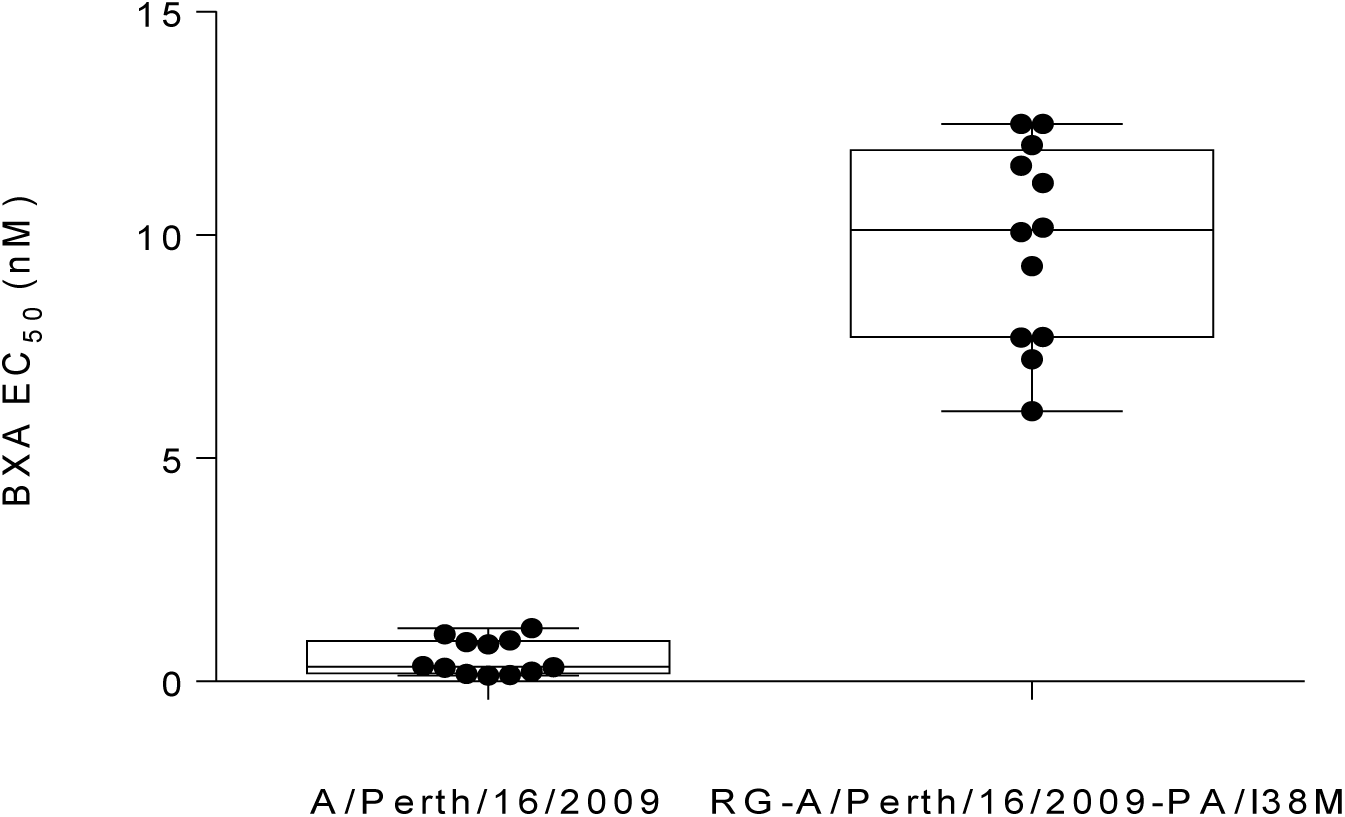
Control influenza virus EC_50_ values for 24-hour FRA used to measure BXA susceptibility. Data was derived from 12 independent experiments, A/Perth/16/2009 had a mean ± standard deviation EC_50_ value of 0.5 ± 0.4 nM and the RG-A/Perth/16/2009-PA/I38M virus had an EC_50_ of 8.5 ± 3.8 nM.

### 3.3 Susceptibility of Circulating Influenza Viruses to BXA

BXA EC_50_ values were obtained for influenza viruses circulating in the Asia-Pacific region between the years 2012 and 2018 (Figure 2). The mean BXA EC_50_ of A(H1N1)pdm09 viruses (n=89) was 0.7 ± 0.5 nM, with a range of 0.1 to 2.1 nM, while n=88 A(H3N2) viruses had a mean EC_50_ of 1.2 ± 0.6 nM with a range of 0.1 to 2.4 nM. The mean EC_50_ value of A(H3N2) viruses were significantly higher than that of A(H1N1)pdm09 viruses (P<0.0001) based on an unpaired student’s T test. B/Victoria lineage viruses (n=53) had a mean EC_50_ of 7.2 ± 3.5 nM, with a range of 0.7 to 14.8 nM, while B/Yamagata lineage viruses (N=56) had a mean value of 5.8 ± 4.5 nM and the range of values of 1.8 nM to 15.5 nM, respectively. Based on a student’s T test there was no significant difference between the mean EC_50_ of influenza B viruses from the two lineages. Taken together the mean EC_50_ of all influenza A viruses (1.0 ± 0.6 nM) was approximately 6-fold lower than that of influenza B viruses (6.6 ± 4.1 nM). This difference was statistically significant (P<0.0001) based on an unpaired student’s T test.

**Figure 2.**
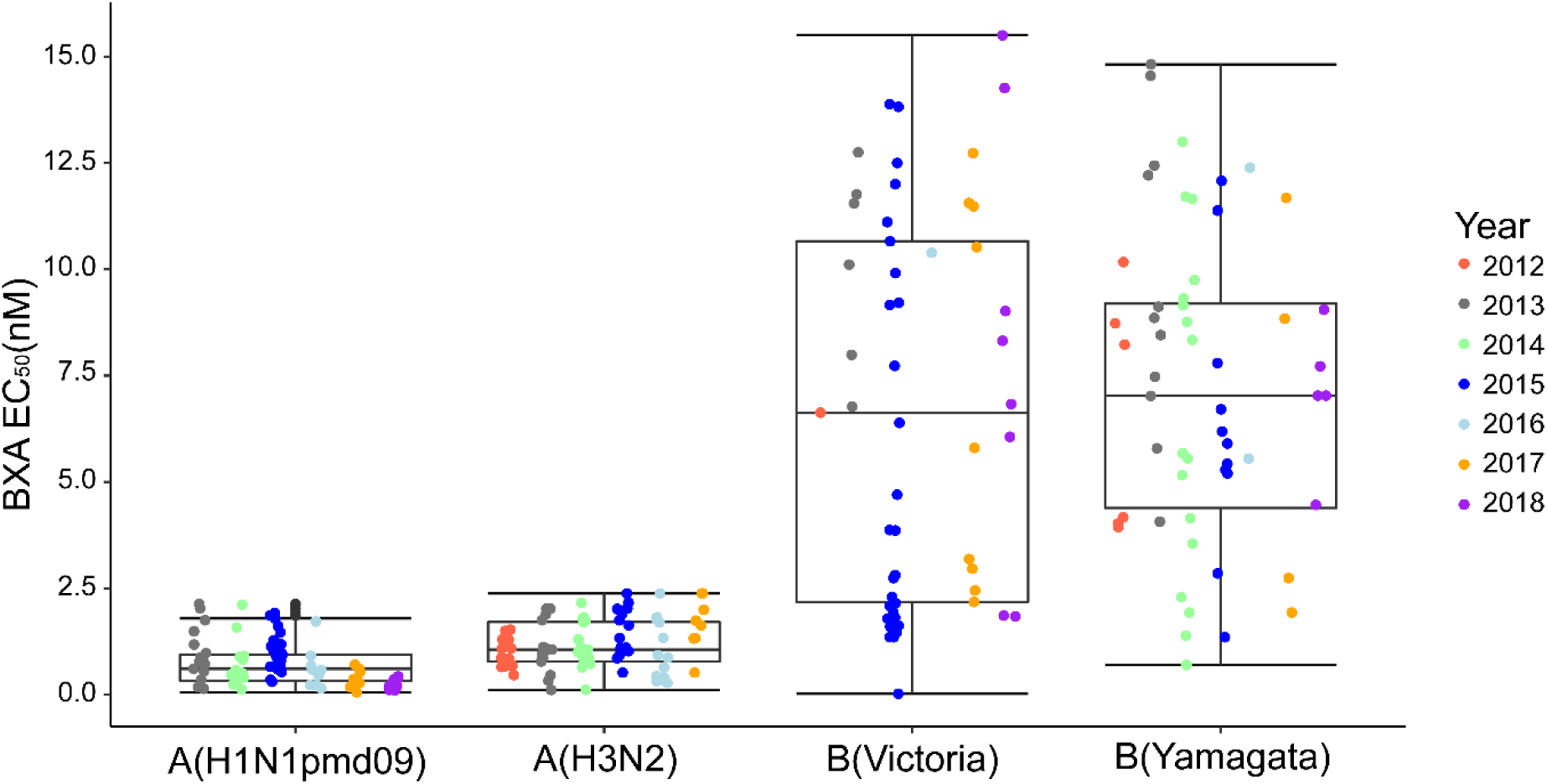
BXA Susceptibility of seasonal influenza viruses circulating in the Asia Pacific region between 2012 and 2018. Influenza A(H1N1pdm09) (n=89), A(H3N2) (n=88) and B(Victoria Lineage) (n=53) and B(Yamagata Lineage) (n=56) were tested in a 24 hour FRA in MDCK-SIAT cells. The EC_50_ was determined using the percentage inhibition of FFU compared to a no-drug but infected virus control well. The mean ± SD EC_50_ values for A(H1N1pdm09), A(H3N2), B/Victoria lineage and B/Yamagata lineage viruses are 0.7 ± 0.5 nM, 1.2 ± 0.6 nM, 7.2 ± 3.5 nM and 5.8 ± 4.5 nM, respectively, and are grouped based on virus subtype/lineage and the year of circulation. Based on a student’s T test the mean EC_50_ of all influenza B viruses was significantly higher than that of influenza A viruses (P<0.0001).

### 3.4 Susceptibility of NAI Resistant Viruses to BXA

Eleven viruses with neuraminidase AA substitutions that confer reduced susceptibility to oseltamivir, zanamivir, peramivir or laninamivir were screened for susceptibility to BXA using the FRA (Table 1). All viruses tested had EC_50_ values within the expected range for influenza A (0.1 to 2.4 nM) and influenza B (0.7 to 15.5 nM). These data demonstrate that BXA is active against influenza strains which have reduced susceptibility to NAIs.

**Table 1.**
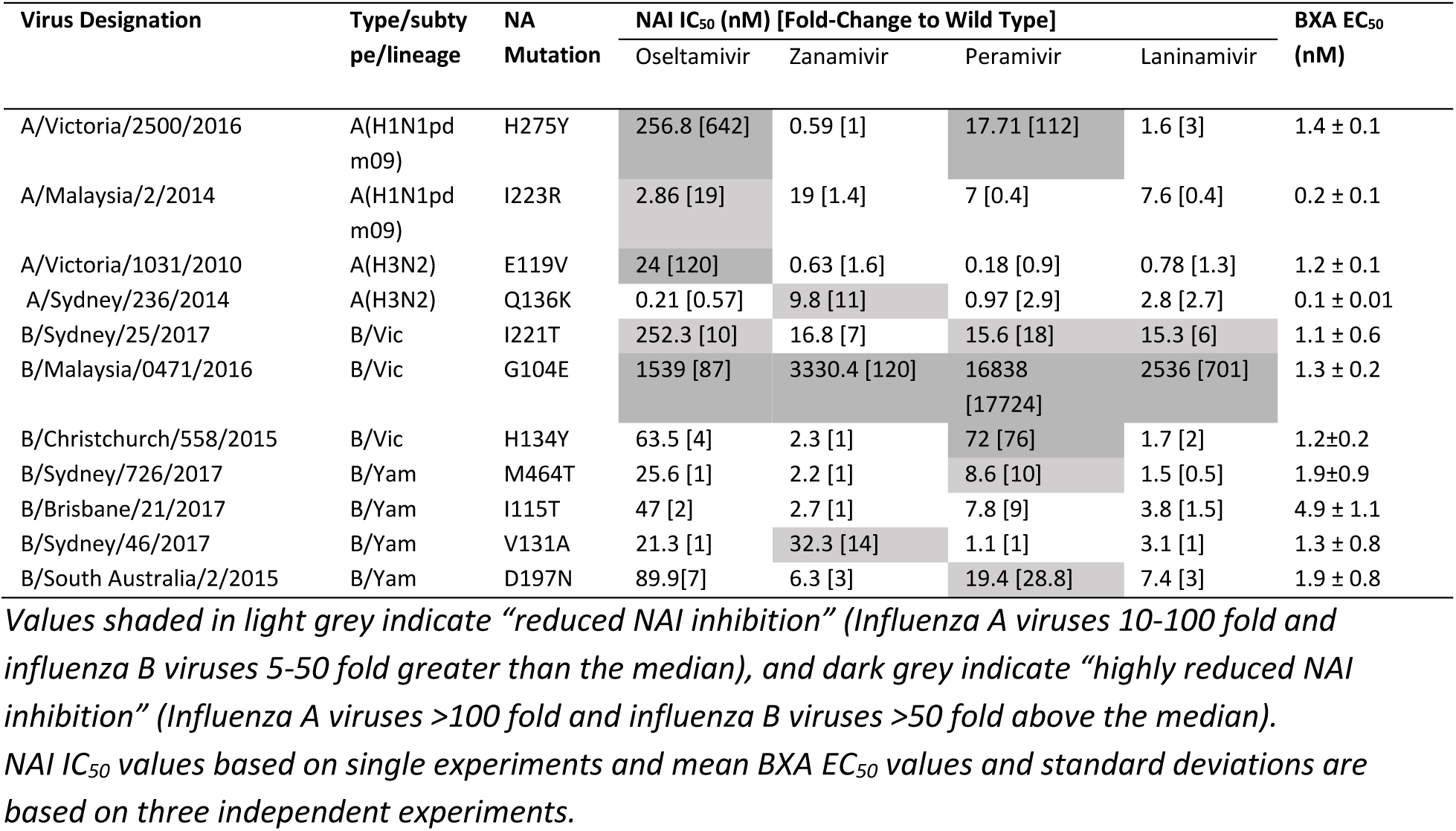
Susceptibility of NAI resistant viruses to BXA.

| Virus Designation | Type/subtype/lineage | NA Mutation | NAI IC<sub>50</sub> (nM) [Fold-Change to Wild Type] |  |  |  | BXA EC<sub>50</sub> (nM) |
| --- | --- | --- | --- | --- | --- | --- | --- |
|  |  |  | Oseltamivir | Zanamivir | Peramivir | Laninamivir |  |
| A/Victoria/2500/2016 | A(H1N1pd m09) | H275Y | 256.8 [642] | 0.59 [1] | 17.71 [112] | 1.6 [3] | 1.4 ± 0.1 |
| A/Malaysia/2/2014 | A(H1N1pd m09) | I223R | 2.86 [19] | 19 [1.4] | 7 [0.4] | 7.6 [0.4] | 0.2 ± 0.1 |
| A/Victoria/1031/2010 | A(H3N2) | E119V | 24 [120] | 0.63 [1.6] | 0.18 [0.9] | 0.78 [1.3] | 1.2 ± 0.1 |
| A/Sydney/236/2014 | A(H3N2) | Q136K | 0.21 [0.57] | 9.8 [11] | 0.97 [2.9] | 2.8 [2.7] | 0.1 ± 0.01 |
| B/Sydney/25/2017 | B/Vic | I221T | 252.3 [10] | 16.8 [7] | 15.6 [18] | 15.3 [6] | 1.1 ± 0.6 |
| B/Malaysia/0471/2016 | B/Vic | G104E | 1539 [87] | 3330.4 [120] | 16838 [17724] | 2536 [701] | 1.3 ± 0.2 |
| B/Christchurch/558/2015 | B/Vic | H134Y | 63.5 [4] | 2.3 [1] | 72 [76] | 1.7 [2] | 1.2±0.2 |
| B/Sydney/726/2017 | B/Yam | M464T | 25.6 [1] | 2.2 [1] | 8.6 [10] | 1.5 [0.5] | 1.9±0.9 |
| B/Brisbane/21/2017 | B/Yam | I115T | 47 [2] | 2.7 [1] | 7.8 [9] | 3.8 [1.5] | 4.9 ± 1.1 |
| B/Sydney/46/2017 | B/Yam | V131A | 21.3 [1] | 32.3 [14] | 1.1 [1] | 3.1 [1] | 1.3 ± 0.8 |
| B/South Australia/2/2015 | B/Yam | D197N | 89.9[7] | 6.3 [3] | 19.4 [28.8] | 7.4 [3] | 1.9 ± 0.8 |

Values shaded in light grey indicate “reduced NAI inhibition” (Influenza A viruses 10-100 fold and influenza B viruses 5-50 fold greater than the median), and dark grey indicate “highly reduced NAI inhibition” (Influenza A viruses >100 fold and influenza B viruses >50 fold above the median).
NAI IC<sub>50</sub> values based on single experiments and mean BXA EC<sub>50</sub> values and standard deviations are based on three independent experiments.

### 3.5 Susceptibility and Phenotypic Screening of PA_N_ I38 Amino Acid Substitution to BXA

Substitutions in the PA_N_ /I38T/I38M/I38F have been identified in some influenza viruses in BXM clinical trials from patients following treatment [12]. Therefore we evaluated the BXA susceptibility of viruses with these substitutions in the FRA. The substitutions PA_N_ /I38T, PA_N_ /I38M and PA_N_ /I38F in an A(H1N1)pdm09 virus conferred a 65, 23 and 17-fold increase in EC_50_ compared to corresponding wild type viruses, respectively, while PA_N_ /I38M and PA_N_ /I38F in an A(H3N2) virus both conferred a 16 fold increase in EC_50_ compared to respective wildtype viruses (Table 2). The PA_N_/I38 AA substitutions conferred smaller increases in EC_50_ in an influenza B virus than they did in the influenza A strains, 5-fold for PA_N_ /I38T and 2-fold for PA_N_ /I38M (Table 2). Of the three AA substitutions, PA_N_ /I38T resulted in the greatest EC_50_ from BXA. RG-A/Perth/16/2009-PA/I38T and RG-B/Yamanashi/166/98-PA/I38F could not be rescued.

**Table 2.** BXA susceptibility of reverse genetics derived influenza viruses with I38 PA_N_ amino acid substitutions.

| Virus Designation (Subtype/Lineage) | PA<sub>N</sub> Protein AA Substitution | BXA EC<sub>50</sub> (nM) [Fold Change from wild type] |
| --- | --- | --- |
| A/Perth/261/2009 (A(H1N1)pdm09) | Wild Type | 0.6 ± 0.5 |
|  | I38T | 37 ± 18.2 [65] ** |
|  | I38M | 13.3 ± 7.3 [23] ** |
|  | I38F | 9.6 ± 5.3 [17] ** |
| A/Perth/16/2009 (A(H3N2)) | Wild Type | 0.5 ± 0.4 |
|  | I38M | 8.5 ± 3.8 [16] **** |
|  | I38F | 8.5 ± 3.7 [16] **** |
| B/Yamanashi/166/98 (B/Yamagata Lineage) | Wild Type | 5.3 ± 4.2 |
|  | I38T | 26.3 ± 18.1 [5] <sup>ns</sup> |
|  | I38M | 9.1 ± 4.1 [2] <sup>ns</sup> |

Mean EC<sub>50</sub> values and standard deviations are based on a minimum of six independent experiments. Fold change in EC<sub>50</sub> is compared to wild type virus for each corresponding subtype.
Unpaired students T Test was used to compare mean EC<sub>50</sub> values of variants to corresponding wild type virus, \* P<0.05, \*\* P<0.01, \*\*\*P<0.001, \*\*\*\*P<0.0001, ns P>0.05.

## 4. Discussion

This study aimed to establish a robust and reproducible assay to determine the susceptibility of influenza viruses circulating in the past seven years to the PA_N_ inhibitor drug BXA. In addition, we investigated the impact of substitutions at position I38 of the PA_N_ in contemporary A(H1N1)pdm09, A(H3N2) and influenza B viruses on BXA susceptibility to confirm the ability of the assay to detect such variants. Using the FRA method, BXA was shown to be active against all 286 influenza viruses tested that had circulated in the Asia-Pacific region from 2012 to 2018. The mean EC_50_ values obtained in this study were similar to those described in a smaller *in vitro* study using a BXA plaque reduction assay and 17 Japanese influenza A viruses (0.2 to 1.0 nM) and four influenza B viruses (4.0 to 11.3 nM)[8].

When comparing the BXA EC_50_ values of influenza A viruses with influenza B viruses, two noteworthy observations can be made. Firstly, mean BXA EC_50_ values are approximately 6-fold higher for influenza B viruses than for influenza A viruses. BXA forms a “wing” shaped structure that binds to five key AAs (20, 24, 34, 37 and 38) in a V-shaped conformation within the PA_N_ active site. However, aside from residue I38, which is conserved across both influenza A and B viruses, the other four positions have different residues in influenza A or B viruses (influenza A viruses: A20, Y24, K34 and A37; influenza B viruses: T20, F24, M34, N37 and I38) [12], which is likely to be the reason for the difference in binding and EC_50_ values. Lower susceptibility in influenza B viruses compared to influenza A viruses is also observed for oseltamivir, where IC_50_ values for influenza B viruses are 15–20 fold higher than that of influenza A viruses [16, 17]. This difference in *in vitro* oseltamivir susceptibility translates into an *in vivo* effect, where numerous studies have reported reduced clinical effect of oseltamivir against influenza B infections compared with influenza A infections [18–21]. However based on clinical trial data in patients at high-risk of severe influenza, BXM seems to have a comparable clinical effect against both influenza A and B virus infections [22]. The second observation of note from this study was that the range of BXA EC_50_ values for influenza B viruses was considerably larger than it was for influenza A viruses, which may be due to greater variation amongst framework residues in the influenza B PA_N_ than for influenza A viruses, resulting in subtle impacts on drug binding.

The interaction of PA_N_ /I38 with BXA is present in both influenza A and B viruses and based on BXM clinical trials, this residue appears prone to selection pressure in both virus types. The I38T AA substitution, which confers the largest change in EC_50_, results in the loss of a methyl group present in wild type viruses. The absence of a methyl group reduces van der Waals interactions between BXM and influenza PA_N_. For drug binding, the presence of T38 also requires a rotational change in the PA_N_ that is not necessary in wild type viruses [12]. Although the substitutions I38M and I38F in A(H1N1)pdm09 and A(H3N2) viruses conferred 16 to 25-fold increases in EC_50_, it is important to note that the EC_50_ values for these viruses range from 8.5 to 13.3 nM, which is not substantially higher than the mean EC_50_ of wildtype influenza B viruses (6.6 ± 4.1 nM). Therefore understanding how these *in vitro* findings impact clinical effectiveness and which of the I38 variants are likely to result in reduced clinical effectiveness will be important.

One disadvantage of the FRA is that it may not be suitable for front-line diagnostic laboratories, where molecular-based genotypic assays are more commonly used due to time, equipment and labour constraints. To date, data indicate that PA_N_/I38 AA substitutions are likely to be the most common AA substitutions that confer reduced BXA susceptibility. While the I38 residue appears to be a ‘hot-spot’ for AA substitutions under BXA pressure, there are a small number of other substitutions in the PA_N_ that have also been reported to reduce susceptibility *in vitro* (such as E199G) [12], and it is likely that additional sites will be detected as clinical use of the drug increases.

This study provides information on the baseline susceptibility of a large number of recently circulating influenza viruses across all relevant subtypes and lineages in the Asia-Pacific region. It will be important to continue to test circulating viruses for BXM susceptibility as the antiviral continues to be licensed and used more widely to better understand the molecular determinants of BXA susceptibility, the frequency with which viruses with reduced susceptibilty occur, and whether they have the capacity to transmit within the community in the absence of drug selective pressure.

## Acknowledgments

The Melbourne WHO Collaborating Centre for Reference and Research on Influenza is supported by the Australian Government Department of Health.

DV is supported by contract HHSN272201400006C from the National Institute of Allergy and Infectious Disease, National Institutes of Health, Department of Health and Human Services, USA

